# Extracellular matrix signals promotes actin-dependent mitochondrial elongation and activity

**DOI:** 10.1101/2024.01.22.576703

**Authors:** Priya Gatti, Pritha Mukherjee, Priyanka Dey Talukdar, Wesley Freppel, Joseph Kanou, Laurent Chatel-Chaix, Urmi Chatterji, Marc Germain

## Abstract

Mitochondria are crucial metabolic organelles that are regulated by both intracellular and extracellular cues. The extracellular matrix (ECM) is a key component of the cellular environment that controls cellular behavior and metabolic activity. Here, we determined how ECM signalling regulates mitochondrial structure and activity. To distinguish mitochondrial regulation from the general survival cues generated by the ECM, we used breast cancer-derived spheres (mammospheres) because of their ability to grow in suspension culture in the absence of ECM. Using this system, we demonstrate that the association of mammospheres with the ECM results in dramatic mitochondrial elongation, along with enhanced mitochondrial respiration and ATP production. This remodeling occurs independently of DRP1 activity, but relies on integrin signaling and actin polymerization. Therefore, our findings demonstrate that ECM-driven actin polymerization plays a crucial role in remodeling mitochondrial networks to promote OXPHOS, which represents a vital step for migrating cells to enhance cellular adhesion and facilitate cell growth.

## Introduction

Cells are highly responsive to their microenvironment, with cells grown in different environments exhibiting distinct phenotypic and functional characteristics (Baharvand et al., 2006, Baker et al., 2010). Factors such as the extracellular matrix (ECM) and nutrient availability, shape cellular behavior and determine cellular fate (Hynes, 2009, Mierke, 2019). In this context, the ECM provides structural support and biochemical cues to cells, controlling their adhesion, migration, proliferation, and differentiation (Dzobo and Dandara, 2023). The attachment of cells to specific ECM components (collagen, laminin, fibronectin) regulates cell morphology, gene expression, and even stem cell fate (Frantz et al., 2010, Discher et al., 2009, Buxboim et al., 2010, Rais et al., 2023). ECM properties, including stiffness (Romani et al., 2022, Baker et al., 2010), growth factors, nutrients and oxygen levels (Case and Waterman, 2015, Baker et al., 2010) thus have an important incidence on cell behavior.

While the ECM alters cellular responses, mitochondria play a crucial role in the regulation of cellular metabolism and apoptosis. This in turn affects cellular behaviours including cell differentiation and stem cell maintenance (Kasahara and Scorrano, 2014, Liesa and Shirihai, 2013). Mitochondrial activity is regulated through changes in mitochondrial structure, including changes in their length and connectivity (Ren et al., 2020). For example, mitochondria elongate in response to amino acid starvation to sustain cellular ATP levels (Gomes et al., 2011, Patten et al., 2014, Thomas et al., 2018, Endo et al., 2020, Rambold et al., 2011). These alterations in mitochondrial structure are controlled by the balance between mitochondrial fusion and fission, which are both dependent on large GTPases of the dynamin family. Mitochondrial fusion requires mitofusins (MFN1/2, outer membrane) and optic atrophy 1 (OPA1, inner membrane) (Chen et al., 2007, Chen et al., 2010, Song et al., 2009, Ban et al., 2017, Cao et al., 2017, Qi et al., 2016), while mitochondrial fission is regulated by Dynamin-related protein 1 (DRP1) (Chakrabarti et al., 2018). Maintaining the balance between mitochondrial fusion and fission is crucial for content exchange (Chen et al., 2010, Dong et al., 2022, Adebayo et al., 2021), mitochondrial DNA (mtDNA) integrity (Sabouny and Shutt, 2021, Silva Ramos et al., 2019), removal of damaged mitochondria (Onishi et al., 2021) and ATP production (Yao et al., 2019, Yang et al., 2023, Hofmann et al., 2023).

Interestingly, recent studies have uncovered an intricate relationship between mitochondrial energy metabolism and the ECM. For example, downregulation of mtDNA-encoded respiratory chain subunits leads to altered ECM composition and stiffness (Bubb et al., 2021). In return, ECM stiffness affects mitochondrial length with softer ECM favoring shorter mitochondria (Romani et al., 2022), and stiffer ECM inhibiting DRP1-dependent mitochondrial fission (Chen et al., 2021). This impacts oxidative phosphorylation (OXPHOS), glutamine metabolism (Papalazarou et al., 2020) and glycolysis-related gene expression (Morris et al., 2016). As ECM stiffness depends on its composition and changes in response to the microenvironment (Yanes and Rainero, 2022), microenvironment alteration can significantly impact cell function, including mitochondrial activity.

The complex interplay between cells and their environment is a key feature of tumours, where cancer cells alter and respond to the tumour microenvironment to promote their survival and immune suppression. A small subset of cancer cells, known as tumour-initiating cells (TICs) or cancer stem cells (CSCs), have stemness features such as self-renewal, clonal proliferation, regeneration, metastasis, and drug resistance (Aponte and Caicedo, 2017, Baharvand et al., 2006). These cells are thought to generate the bulk of the cancer cells within the tumour. Importantly, these cells can be isolated through their ability to form spheres when grown in suspension. Nevertheless, the characteristics of CSCs, including their metabolic profile and mitochondrial status remains debated.

Here, we demonstrate that breast cancer cells grown as spheres (mammospheres) alter their mitochondrial structure and metabolic state in an actin-dependent manner upon attachment to the ECM. Specifically, mammospheres exhibit fragmented mitochondria and lower ATP production, but dramatically elongate their mitochondria and increase OXPHOS upon ECM attachment. This remodeling occurs independently of DRP1 activity, but relies on integrin signaling and actin polymerization as interfering with either process prevents it. Therefore, our findings demonstrate that ECM-driven actin polymerization plays a crucial role in remodeling mitochondrial networks to promote OXPHOS, which represents a vital step for migrating cells to enhance cellular adhesion and facilitate cell growth.

## Results

### Breast cancer cells grown as mammospheres have a distinct mitochondrial structure

Some cancer cells can be cultured both as monolayer and suspension cultures, where they adopt distinct metabolic and proliferative phenotypes. For example, a subset of cells within a cancer cell population can grow in suspension as spheres with stem-like properties. These CSCs or TICs have metabolic properties that are distinct from their parent cells, although their study have yielded conflicting results (Andrzejewski et al., 2017, Endo et al., 2020, Pasto et al., 2014). As ECM stiffness alters mitochondrial structure and CSCs are known to be metabolically flexible, we tested the possibility that the interaction between spheres and the ECM controls their metabolic state.

To address this question, we used the aggressive metastatic breast cancer cells MDA-MB-231 (triple negative) that can be grown as an adherent monolayer (AM) and as spheres in suspension (mammospheres, MS)(Wang et al., 2014). Compared to their counterpart grown as a monolayer, MDA-MB-231 mammospheres had decreased ATP and lactate levels (Figure 1A-B), consistent with these cells being less metabolically active (Chen et al., 2021b, Cheung and Rando, 2013). As mitochondrial structure is a key determinant of mitochondrial function, we then determined whether the metabolic differences we observed were associated with changes in mitochondrial structure. MDA-MB-231 cells grown as a monolayer or mammospheres were labelled with the mitochondrial outer membrane protein TOM20 and imaged by confocal microscopy. Mitochondrial structures were then manually quantified as long, intermediate, or fragmented (see methods). MDA-MB-231 cells grown as an adherent monolayer displayed a mixed mitochondrial phenotype, with the majority of mitochondria exhibiting an intermediate phenotype (Figure 1C-D). In contrast, mammospheres showed highly fragmented mitochondria (Figure 1C-D), correlating with the decrease in ATP levels.

**Figure 1.**
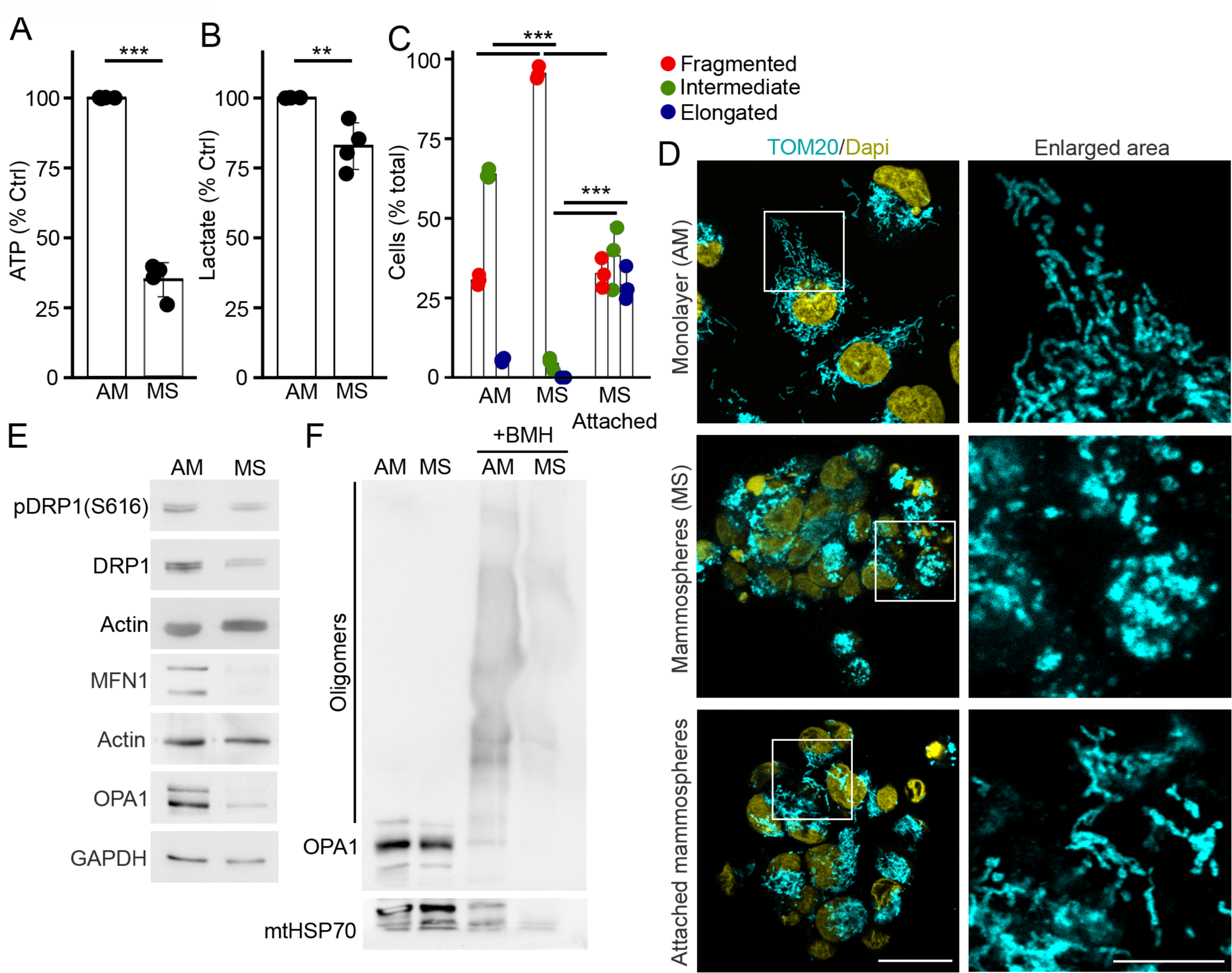
Mammospheres have decreased metabolism and fragmented mitochondria. (A-B) Measure of ATP (A) and lactate (B) levels in MDA-MB-231 cells grown as an adherent monolayer (AM) or as mammospheres in suspension (MS). Each point represents an individual experiment. Bars show the average of 4 independent experiments ± SD. *** p<0.001, ** p<0.01. Two-sided t-test (C-D) Mitochondrial fragmentation in mammospheres in suspension. MDA-MB-231 were grown as AM, MS or MS attached for 6 hours on glass coverslips, and their mitochondria were marked with an antibody against TOM20 (mitochondria, cyan; with nuclei stained with DAPI, yellow). Quantification of 3 independent experiments is shown in (C), with each point represents an individual experiment. Bars show the average ± SD. *** p<0.001. Two-way ANOVA. Representative images are shown in (D). Scale bar 10 µm. (E-F) Western blot showing the expression of mitochondrial dynamics GTPases (E) or OPA1 oligomerization (F). Actin (E) or mtHSP70 (F, lower band – the upper band represents leftover OPA1 signal) were used as loading controls.

To gain further insights into the status of mitochondrial dynamics, we then quantified the expression of fission and fusion proteins. Consistent with the observed mitochondrial fragmentation in mammospheres, we noted a significant decrease in the expression of the fusion proteins MFN1 and OPA1 in these cells (Figure 1E). This was accompanied by a decrease in OPA1 oligomerization, which is required for OPA1 fusion activity (Figure 1F). Nevertheless, mammospheres also exhibited low levels of the fission protein DRP1, both in its active phosphorylated state (pS616) and in terms of total protein levels (Figure 1E), suggesting an overall suppression of mitochondrial dynamics in these cells.

### ECM attachment drives mitochondrial elongation

The above data is consistent with mammospheres exhibiting a mitochondrial profile that is distinct from adherent monolayer cells. As one difference in culture conditions between adherent cells and mammospheres is the absence of attachment signals in the latter, we tested whether attachment alters mitochondrial dynamics in mammospheres. We thus attached mammospheres onto coverslips while maintaining other culture parameters identical, and measured their mitochondrial structure. Compared to mammospheres in suspension, attached mammospheres exhibited elongated mitochondrial structures (Figure 1C-D). A similar pattern was also observed in the less invasive MCF-7 cells (estrogen receptor-positive) (Supp Figure 1). Overall, these results indicate that attachment signals promote mitochondrial elongation in mammospheres.

While mammospheres were attached directly onto glass coverslips in this experiment, cells normally interact with components of the ECM that promote their adhesion. We thus tested the impact of adhesion molecules on mitochondrial elongation using three conditions: 1) Direct attachment to an uncoated plate without ECM, 2) Attachment to a plate coated with the cationic polymer Poly-D-Lysine (PDL) as an ECM control (Lu et al., 2009) and 3) Attachment to a plate coated with the ECM component fibronectin. Mammospheres were transferred to these plates and mitochondrial length quatified over a 6-hour time course. While it took 6 hours of attachment to reach 50% of cells with elongated mitochondria in the absence of substrate, the presence of PDL reduced this time to 2 hours (Figure 2A). Importantly, the presence of the ECM adhesion molecule fibronectin significantly expedited mitochondrial elongation, with most cells showing elongated mitochondria within one hour of attachment (Figure 2A). Similar results were also observed in MCF-7 mammospheres upon attachment to fibronectin and PDL (Figure 2B). Attachment to the ECM thus triggers mitochondrial elongation in mammospheres.

**Figure 2.**
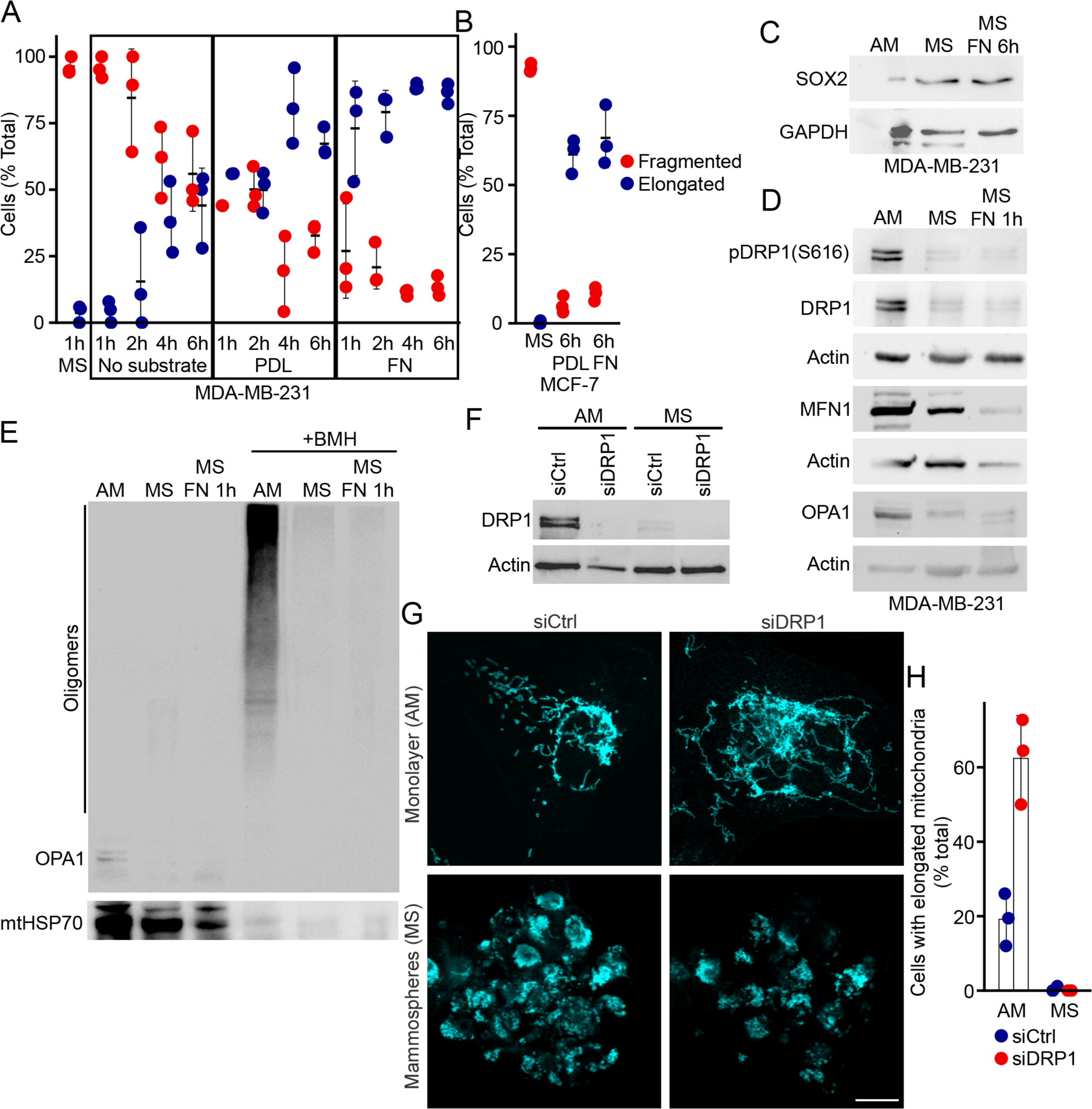
Cell attachment to the ECM promotes mitochondrial elongation. (A-B) Mitochondrial elongation in response to cellular attachment to ECM substrates. MDA-MB-231 (A) and MCF7 (B) were spun down on coverslips either not coated or coated with poly-D-lysin (PDL) or fibronectin (FN) and incubated for the indicated times. mitochondria were marked with an antibody against TOM20 and mitochondrial length quantified in 3 independent experiments. Each point represents an individual experiment. Bars show the average ± SD. (C-D) Western blot showing the expression of the stem cell marker SOX2 (C) and mitochondrial dynamics GTPases in MDA-MB-231 cells grown as AM, MS or MS attached for 6 hours on fibronectin. Actin (C) or GAPDH (D) were used as loading controls. Western blot showing the lack of OPA1 oligomerization in MS cells in the absence or the presence of fibronectin. mtHSP70 was used as a loading control. (F-H) DRP1 knockdown does not affect mitochondrial structure in MDA-MB-231 mammospheres. (F) Western blot showing the knockdown of DRP1 in MDA-MB-231 cells. (G) Representative images showing mitochondrial structure (marked with an antibody against TOM20) in MDA-MB-231 cells knocked down for DRP1. Scale bar 10 µm. Mitochondrial length was quantified in 3 independent experiments (H) Each point represents an individual experiment. Bars show the average ± SD.

### ECM attachment does not alter mitochondrial dynamins

To define the mechanism through which attachment to the ECM triggers mitochondrial elongation, we first verified that this is not the result of changes in their stem-like properties. For this, we measured the expression of the stem cell marker SOX2, which is expressed at high levels in mammospheres (Liu et al., 2018). Consistent with this, SOX2 levels were high in mammospheres compared to adherent monolayer cells and, importantly, SOX2 expression was maintained upon attachment (Figure 2C), indicating that these attachment conditions do not alter the stem-like characteristics of mammospheres.

The dynamins MFN1/2 and OPA1 play a key role in the regulation of mitochondrial fusion. Nevertheless, the expression level of MFN1 and OPA1 did not change upon attachment to fibronectin (Figure 2D). Similarly, attachment to fibronectin did not promote OPA1 oligomerisation (Figure 2E), suggesting that ECM-dependent mitochondrial elongation is not caused by changes in MFN1 or OPA1 expression. Nevertheless, some processing of OPA1 to shorter isoforms did occur (Figure 2D), which has been linked to both mitochondrial fission and facilitated fusion of the inner mitochondrial membrane (Ban et al., 2017, Wang et al., 2021).

It was recently suggested that changes in mitochondrial length triggered by alterations in ECM stiffness in adherent cells is dependent on the mitochondrial fission protein DRP1 (Romani et al., 2022). As mammospheres grown in suspension versus attached could be functionally similar to a change from soft to stiff ECM, we then addressed the role of DRP1 in ECM-dependent mitochondrial elongation. Total DRP1 and its active phosphorylated version (pS616) remained very low following attachment to fibronectin (Figure 2D), and most cells had elongated mitochondria (Figure 2A). While this suggested that DRP1 is not involved in ECM-dependent mitochondrial elongation, we sought to confirm this by knocking down DRP1 using siRNA. Knockdown of DRP1 in adherent monolayer cells caused an elongated mitochondrial phenotype (Figure 2F-H), consistent with its role in promoting mitochondrial fission. In contrast, silencing of DRP1 had no effect on mitochondrial structure in mammospheres (Figure 2F-H), with most cells still exhibiting fragmented mitochondria. This suggested that ECM-triggered mitochondrial elongation in mammospheres operates through a DRP1-independent mechanism, implying the involvement of factors other than expression levels of fission/fusion dynamins in promoting mitochondrial elongation.

### Actin polymerization drives mitochondrial elongation

Given that rearrangement of the actin cytoskeleton is a key event downstream of ECM attachment and that actin is required for mitochondrial fission (Chakrabarti et al., 2018, Hatch et al., 2014, Korobova et al., 2013, Li et al., 2015), we then tested whether actin regulates mitochondrial elongation in attached mammospheres. We first labeled MDA-MB-231 mammospheres in suspension with the filamentous actin (F-actin) stain phalloidin and the mitochondrial marker TOM20. Most mammospheres showed a faint/absent signal for F-actin, except for a small subset of cells displaying strong F-actin signal and elongated mitochondria compared to F-actin low cells (Figure 3A-B). In contrast, attached mammospheres exhibited a prominent F-actin signal (Figure 3C). Specifically, PDL and fibronectin treatments resulted in rapid cell spreading accompanied by well-developed actin structures (Figure 3C) that correlated with the rapid changes in mitochondrial network that occur in these cells (Figure 2A). As MCF7 mammospheres did contain some cells with intermediate mitochondria (Sup. Figure 1), we also assessed actin status in these cells. In contrast to MDA mammospheres, where actin was barely detectable in most cells, the majority of MCF7 cells in mammospheres had weak but clearly visible phalloidin signal that correlated with the presence of intermediate mitochondria (Figure 3A-B, images of MDA and MCF7 cells were taken together using the same settings, allowing comparisons).

**Figure 3.**
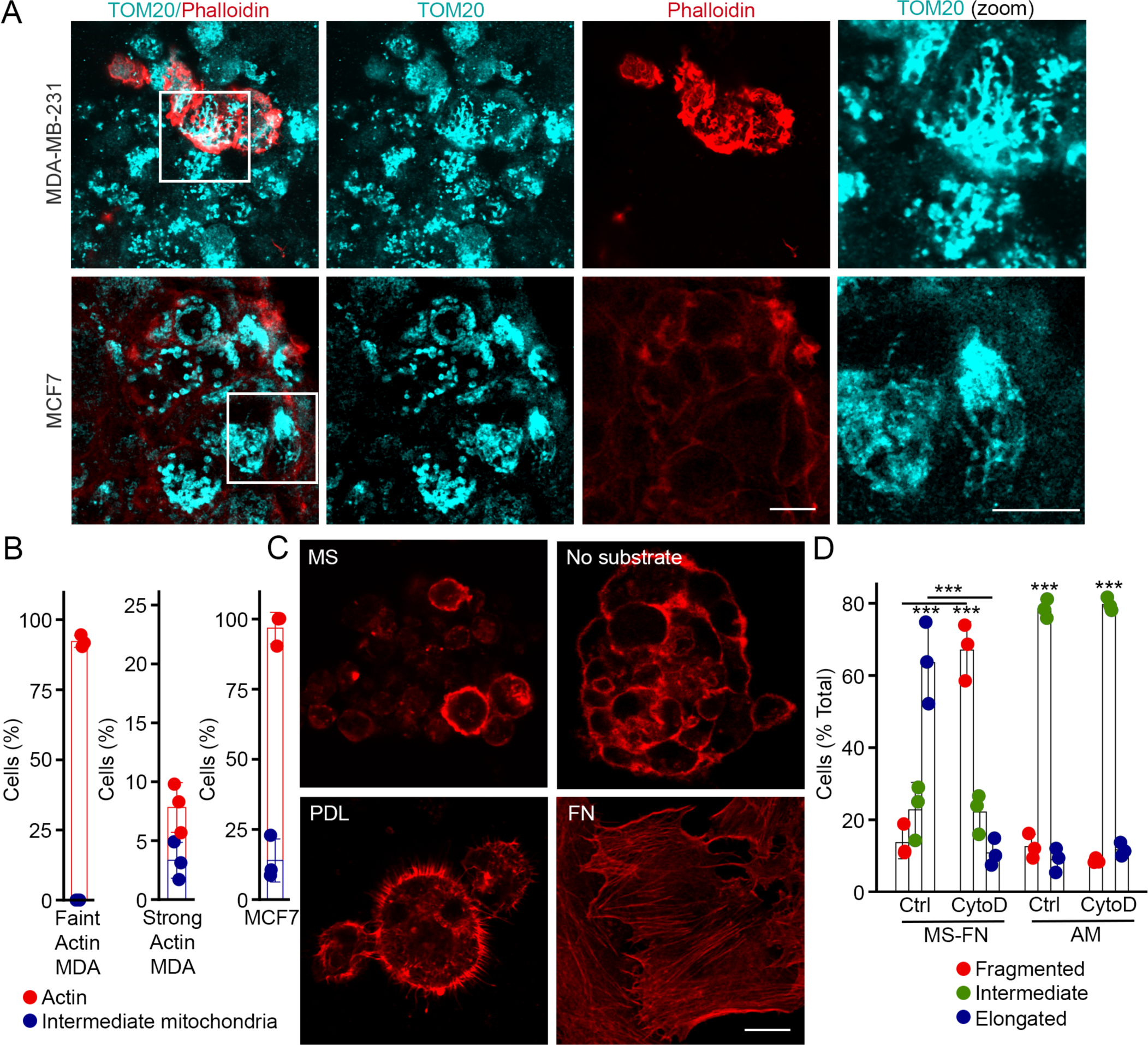
Actin polymerization drives mitochondrial elongation. (A-B) F-Actin staining in MDA-MB-231 and MCF7 mammospheres in suspension. Cells were marked for F-Actin (phalloidin, red) and mitochondria (TOM20, cyan). Scale bar 10 µm. The number of cells with intermediate mitochondria was quantified in MDA-MB-231 cells with faint (B, left) or strong (B, middle) actin staining. As most MCF7 cells had similar F-Actin staining, they were all grouped together for mitochondrial length analysis (B, right). Each point represents an individual experiment. Bars show the average of 3 independent experiments ± SD. (C) Representative images showing F-Actin structures (labelled with phalloidin) present in MDA-MB-231 mammospheres under different attachment conditions. Scale bar 10 µm. (D) Actin depolymerization prevents mitochondrial elongation in mammospheres. MDA-MB-231 cells were grown as an adherent monolayer (AM) or as mammospheres attached for 1 hour on fibronectin (MS-FN) in the absence or the presence of the actin depolymerizing agent CytoD (0.1 μg/ml) and mitochondria were analysed as in (B). Each point represents an individual experiment. Bars show the average of 3 independent experiments ± SD. *** p<0.001.

Given the correlation between actin polymerization and mitochondrial elongation, we then inhibited actin polymerization using Cytochalasin D (CytoD) and determine its effect on mitochondrial networks. Inhibiting actin polymerization efficiently prevented fibronectin-activated mitochondrial elongation (Figure 3D) in attached mammospheres, consistent with an important role for actin in this process. However, CytoD did not significantly affect cells in adherent monolayers (Figure 3D), suggesting that actin is especially required during the elongation of the network (as opposed to steady-state maintenance).

### Mitochondrial Elongation is dependent on signalling downstream of integrins

Cellular attachment to fibronectin is mediated by integrins, which subsequently activates Focal Adhesion Kinase (FAK) and Src, ultimately promoting actin polymerization (Sup. Figure 2A). Consistent with this, while levels of phophorylated FAK (pY397-FAK) were undetectable in mammospheres in suspension, its levels dramatically increased following one hour of attachment to fibronectin (Figure 4A). To ascertain whether FAK activation is required for mitochondrial elongation following cell attachment, we inhibited FAK using the small molecule inhibitor PF-573228 (PF). Consistent with the role of FAK in cellular attachment and actin polymerization, FAK inhibition led to a significant decrease in cell adhesion to fibronectin coated plates (Figure 4B) and impaired their ability to spread and form lamellipodia-like actin features (Figure 4C).

**Figure 4.**
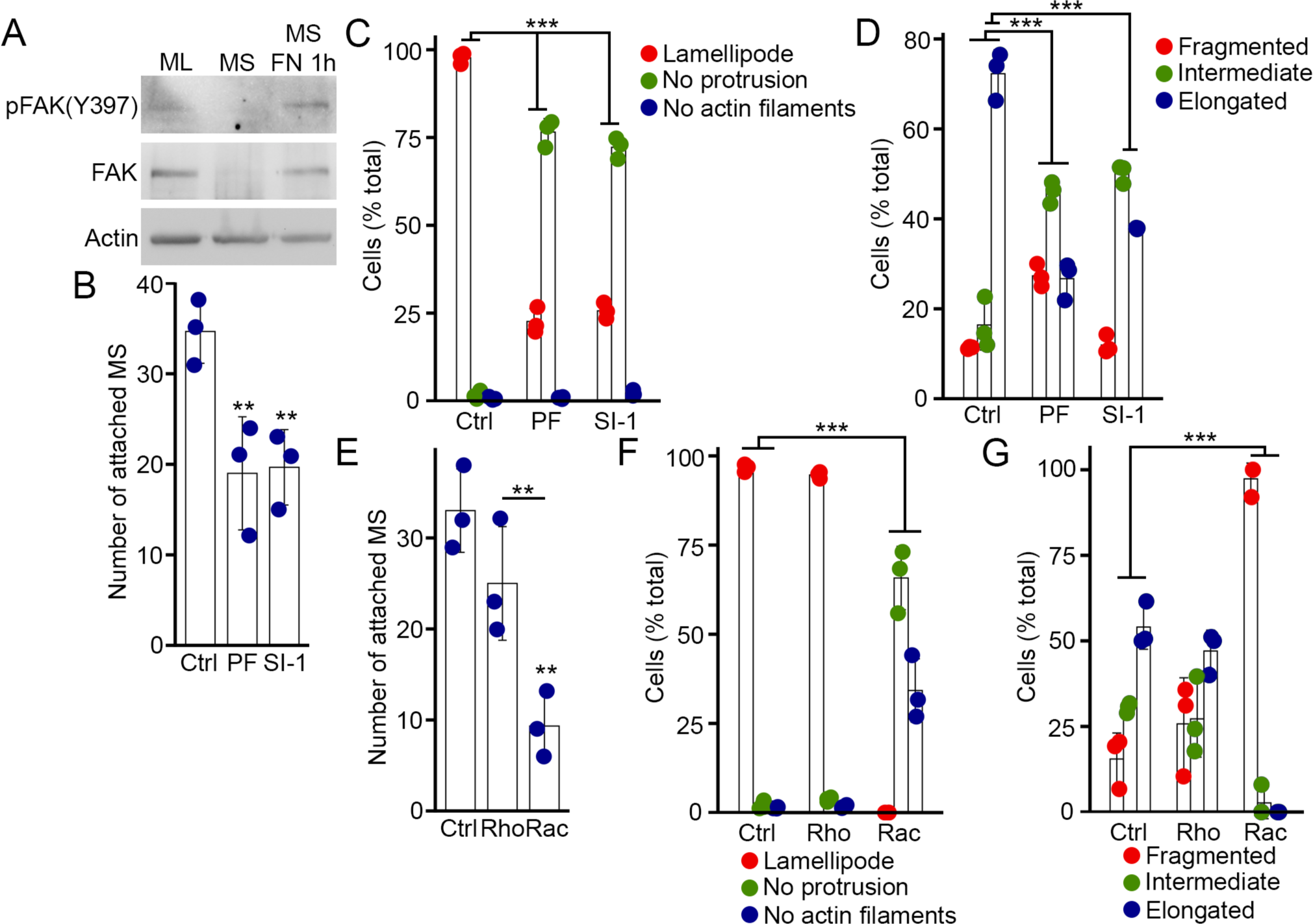
Inhibition of the signalling pathway downstream of integrins prevents mitochondrial elongation. (A) Western blot showing FAK activation (pFAK-Y397) in response to attachment to fibronectin. Actin was used as a loading control. (B-D) Effect of the FAK inhibitor PF-573228 (PF, 10 μM) and the Src inhibitor Src Inhibitor-1 (SI-1, 10 μM) on mammosphere attachment to fibronectin-coated plates (1 hour) (B) formation of F-actin structures (C) and mitochondrial structure (D). Each point represents an independent experiment. Bars show the average ± SD. *** p<0.001. (E-G) Effect of Rac1 (Rac Inhibitor III; 2 μM) and Rho (Rho Inhibitor I, 10μM) inhibition on mammosphere attachment to fibronectin-coated plates (1 hour) (E), formation of F-actin structures (F) and mitochondrial structure (G). Each point represents an independent experiment. Bars show the average ± SD. *** p<0.001. One-way ANOVA (B, E), Two-way ANOVA (C, D, F, G).

Importantly, FAK inhibition caused a profound impairment of mitochondrial elongation in the cells that did attach to fibronectin-coated plates (Figure 4C-D), consistent with this pathway being required for fibronectin-driven mitochondrial elongation. Src is a tyrosine kinase found within the integrin adhesion complex that facilitates FAK activation and promotes maximal adhesion induced by FAK activation (Calalb et al., 1995). In fact, inhibition of Src in attached mammospheres using Src inhibitor-1 (SI-1) inhibited attachment and actin polymerization similarly to FAK inhibition (Figure 4B-C). More importantly, Src inhibition significantly inhibited mitochondrial elongation (Figure 4D), further supporting a role for integrin signalling in fibronectin-driven mitochondrial elongation.

Rho GTPases such as RhoA, Cdc42, and Rac1 are the key signalling proteins that act downstream of FAK/Src to promote actin cytoskeleton organization. Among these, Rac1 regulates the formation of lamellipodia-like actin structures (Horton et al., 2015, Price et al., 1998) and its expression is elevated in cancer stem cells (Ko et al., 2014, Sundberg et al., 2003). Consistent with this, inhibition of Rac1 (with Rac Inhibitor III) reduced attachment to fibronectin and actin rearrangements in the cells that did attach (Figure 4E-F, Sup. Figure 2B). Rac1 inhibition also completely prevented mitochondrial elongation upon attachment (Figure 4G), further supporting a crucial role for actin in promoting mitochondrial elongation. On the other hand, RhoA inhibition (with Rho Inhibitor I) did not significantly affect attachment or mitochondrial structure (Figure 4E-G), suggesting that actin reorganization and mitochondrial elongation is mainly Rac-dependent in our experimental setting.

While our results indicate that a signalling pathway downstream of integrins is required for mitochondrial elongation upon attachment to fibronectin, the partial inhibition of cell attachment caused by the inhibition of this pathway could potentially influence this conclusion. As an alternative, we thus directly activated Rho GTPases in mammospheres in suspension culture using a small molecule known to activate RhoA, Cdc42, and Rac1 (Rho/Rac/Cdc42 Activator I). Following activation, we visualized the cells by labeling mitochondria (with TOM20) and actin (with phalloidin). While control mammospheres contained only a few cells with strong actin staining, the majority of cells treated with the activator displayed an intense F-actin signal (Figure 5A-B). More importantly, this was associated with an increase in the number of cells displaying elongated mitochondria (Figure 5A and C), further demonstrating that mitochondrial elongation occurs downstream of Rho family-driven actin polymerization.

**Figure 5.**
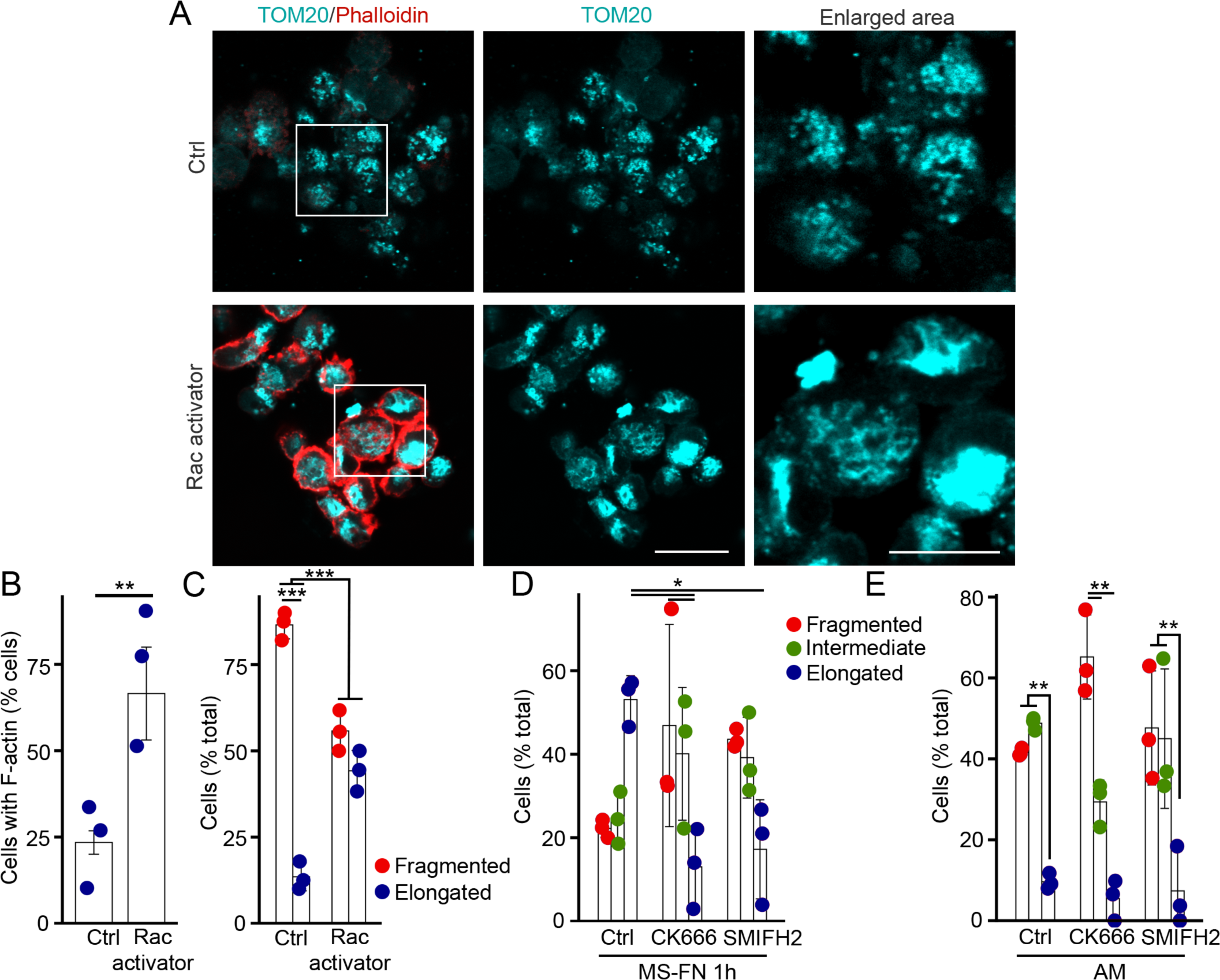
Actin polymerization promotes mitochondrial elongation. (A-C) Actin polymerization and mitochondria in mammospheres treated with a Rac activator (Rho/Rac/Cdc42 Activator I, 10 μM, 1 hour). (A) Representative images of F-actin (phalloidin, red) and mitochondria (TOM20, cyan) in MDA-MB-231 mammospheres. Scale bar 10 µm. The quantification of 3 independent experiments is shown in (B) for F-actin and (C) for mitochondrial length. Each point represents an individual experiment. Bars show the average ± SD. *** p<0.001. two-sided t-test (B), Two-way ANOVA (C). (D-E) Disruption of mitochondrial elongation in mammospheres treated with the Arp2/3 inhibitor CK666 (10 μM, 1 hour) or the formin inhibitor SMIFH2 (2 µm, 1 hour). Mitochondrial length was quantified in mammospheres attached to fibronectin (1 hour) (D) and attached monolayer cells (E). Each point represents an individual experiment. Bars show the average ± SD of 3 independent experiments. ** p<0.01, * p<0.05. Two-way ANOVA.

Rac1 promotes lamellipodia formation by activating the Wiskott-Aldrich syndrome family of proteins (WASP, N-WASP, and WAVE1/2) that, in turn, activates actin-polymerizing proteins including the Arp2/3 complex and formins. Consistent with this, inhibition of Arp2/3 (using CK666) or formins (using SMIFH2) significantly reduced mitochondrial elongation in attached mammospheres (Figure 5D). On the other hand, the effect of the inhibitors was limited in adherent monolayer cells, with only CK666 causing an increase in cells with fragmented mitochondria relative to those with intermediate mitochondria (Figure 5E). While this is in line with the lack of effect of CytoD in these cells (Figure 3D), it suggests that actin plays a more prominent role during widespread changes in mitochondrial structure in response to changes in cellular environment.

### Mitochondrial elongation promotes OXPHOS in mammospheres

Alterations in mitochondrial architecture exert a profound influence on mitochondrial function, with mitochondrial fusion or elongation being vital in preserving mitochondrial function during starvation (Adebayo et al., 2021). We thus investigated whether the observed alterations in mitochondrial dynamics during cellular attachment impact mitochondrial activity. We first determined whether the changes in mitochondrial structure were associated with alterations in the expression of electron transport chain (ETC) proteins that are required for OXPHOS. In contrast to the decrease in ATP production (Figure 1A), mammospheres exhibited higher expression of all ETC components tested relative to their parental cells grown as an attached monolayer (Figure 6A). Interestingly, the expression of ETC components was partially decreased upon attachment to fibronectin (Figure 6A), suggesting a direct relationship between mitochondrial structure and ETC component expression. Nevertheless, mitochondrial membrane potential remained similar across conditions (Figure 6B), suggesting that they remain functional irrespective of their morphology.

**Figure 6.**
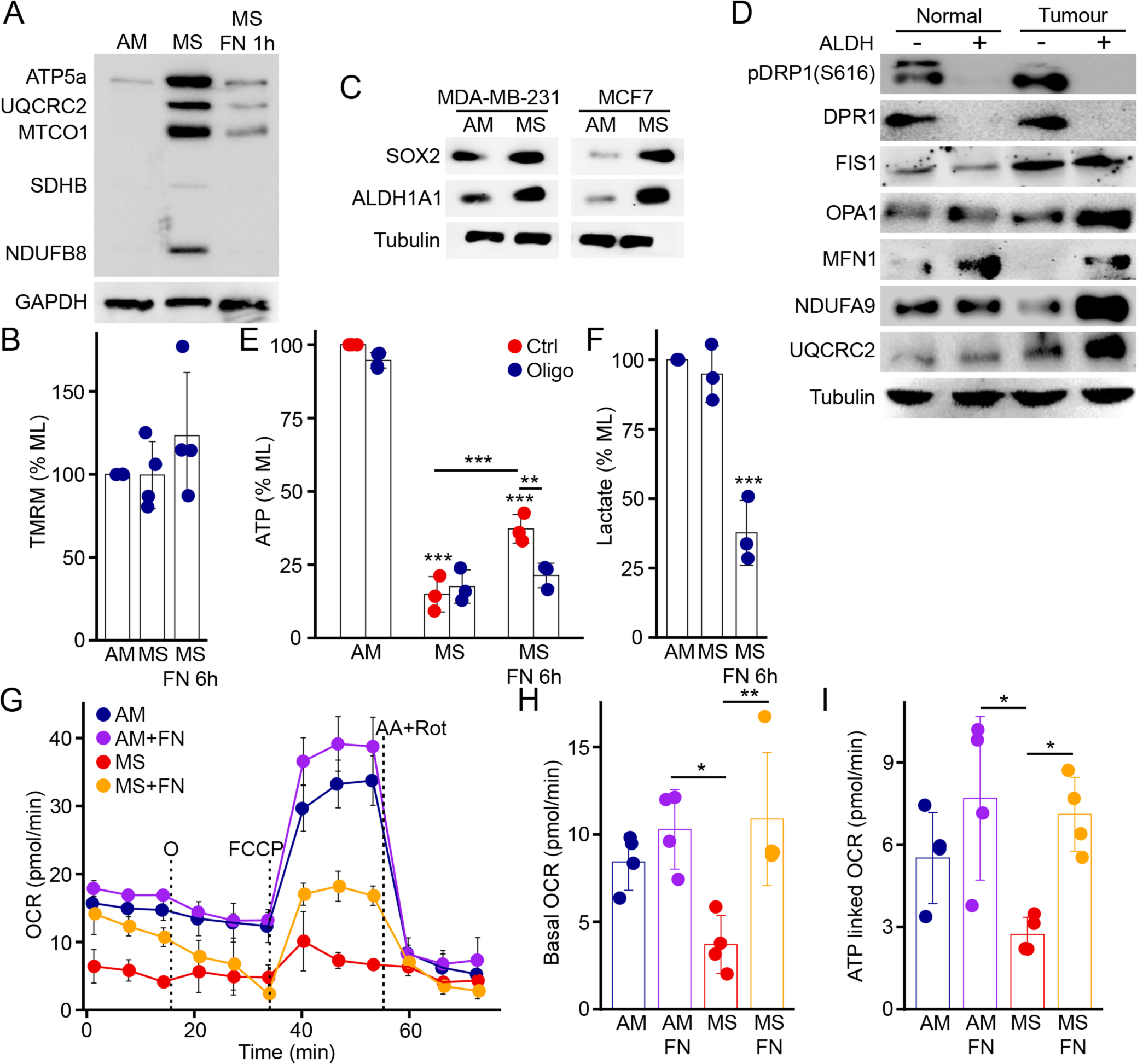
Mitochondrial elongation promotes OXPHOS in mammospheres. (A) Western blot showing the expression levels of ETC components in attached monolayer cells (AM), mammospheres (MS) and mammospheres attached to fibronectin for 1 hour (MS-FN). GAPDH was used as a loading control. (B) Mitochondrial membrane potential measured using TMRM in AM, MS and MS attached to fibronectin for 1 hour (MS-FN) in 3 independent experiments. Each point represents an individual experiment. Bars show the average ± SD. (C) (C) Western blot showing the expression levels of the stem cell markers SOX2 and ALDH1A1 in MDA-MB-231 and MCF7 cells grown as an adherent monolayer or as mammospheres in suspension. β-Tubulin was used as a loading control. (D) Western blot showing the expression levels of mitochondrial proteins in ALDH-negative and positive cells isolated from normal mammary gland and mammary tumour tissues. β-Tubulin was used as a loading control. (E-F) Metabolic changes in MDA-MB-231 mammospheres upon attachment to fibronectin. Total ATP (E) and lactate (F) were measured in 3 independent experiments. Each point represents an individual experiment. Bars show the average ± SD. *** p<0.001, ** p<0.01. one-way ANOVA, two-way ANOVA for (B). (G-I) Changes in Oxygen consumption in mammospheres upon attachment to fibronectin. (G) Representative oxygen consumption rate (OCR) curves. O, oligomycin; AA, antimycin A; Rot, rotenone. Basal OCR (H) and ATP-linked OCR (I) were calculated from 4 independent experiments. Each point represents an individual experiment. Bars show the average ± SD. ** p<0.01, * p<0.05. One-way ANOVA.

As previously noted, mammospheres are known to house a high proportion of CSCs, confirmed by the expression of the stem cell markers SOX-2 and ALDHIA1 in MDA-MB-231 and MCF-7 mammospheres (Figure 6C). We explored the relevance of these findings in an *in vivo* context by isolating ALDH-positive breast CSCs (brCSCs) from mouse mammary tumors. Consistent with the mammosphere data, brCSCs displayed reduced expression of the fission protein DRP1 (both total and pDRP1; Figure 6D). These cells also had increased levels of the mitochondrial fusion proteins OPA1 and MFN1 (Figure 6D), likely reflecting their long-term interaction with the ECM within the tumour environment. Importantly, brCSCs also showed increased expression of the ETC proteins NDUFA9 and UQCRC2, mirroring our *in vitro* data (Figure 6D).

To better understand the consequences of ECM attachment on mitochondrial OXPHOS, we measured bioenergetic parameters (ATP and Lactate) in adherent monolayers, mammospheres, and attached mammospheres (FN-6 hours). Consistent with our previous results (Figure 1A-B), mammospheres exhibited lower ATP levels than adherent monolayer cells (Figure 6E). Importantly, attachment to fibronectin partially rescued ATP levels in mammospheres (Figure 6E), in line with the mitochondrial elongation observed following attachment. We then assessed the mitochondrial contribution to ATP levels using the ATP synthase inhibitor oligomycin. Oligomycin had little effect on adherent monolayers or mammospheres (Figure 6E), indicating limited reliance on mitochondria-derived ATP. In contrast, oligomycin significantly reduced ATP levels in attached mammospheres, suggesting increased mitochondria dependence. This correlated with a significant reduction in lactate levels upon attachment (Figure 6F). This indicates that, following attachment to fibronectin, cells stimulate their mitochondrial metabolism, promoting their dependency on OXPHOS for ATP production.

To further confirm that ECM attachment promotes mitochondrial dependency for energy production, we measured oxygen consumption rates (OCR). Consistent with mammospheres being less metabolically active, they showed lower basal OCR and minimal ATP-linked respiration compared to monolayer cells (Figure 6G-I). Importantly, attached mammospheres significantly increased both basal and ATP-linked OCR, consistent with mitochondrial elongation stimulating mitochondrial activity (Figure 6G-I). Altogether, our results indicate that the actin-driven mitochondrial elongation that occurs upon attachment to fibronectin stimulates mitochondrial respiration in previously less metabolically active mammospheres.

## Discussion

The ECM plays a crucial role in the modulation of cell survival and function. Previous studies have also suggested links between ECM signalling and mitochondria (Bubb et al., 2021, Romani et al., 2022, Visavadiya et al., 2016, Cai et al., 2023, Tian et al., 2023). While these observations could mechanistically link the migration, proliferation, and survival roles of ECM signalling with metabolic needs, the complex links between these events makes it challenging to identify clear causal links in adherent cells requiring ECM signals for their survival. Here, we took advantage of the fact that a subset of cells within a tumour can be grown in suspension in the absence of ECM. These mammospheres have distinct metabolic properties and mitochondrial structure. As mitochondrial dynamics regulates mitochondrial activity, we used these cells to investigate how mitochondrial structure and function are regulated by the ECM. We show that even if mammospheres do not require an ECM for their survival, they modulate their mitochondrial structure and energy metabolism in response to ECM components. We also uncovered a key role for actin polymerization in this process.

Some studies have reported that the ECM regulates mitochondrial morphology in response to ECM stiffness. Stiff ECM has been found to block mitochondrial fission (Chen et al., 2021a), while soft ECM promotes it (Romani et al., 2022). Stiff ECM activates β1-integrin/PINCH1-kindlin-2 signaling, which inhibits DRP1 activity, thereby promoting mitochondrial elongation (Chen et al., 2021a). In contrast, soft ECM activates DRP1, leading to mitochondrial fission and metabolic alterations through the modulation of mitochondrial ROS and antioxidants (Romani et al., 2022). In contrast, the mitochondrial elongation we observed upon mammospheres attachment is unlikely to involve DRP1. Despite mammospheres in suspension (akin to soft ECM) having fragmented mitochondria and becoming elongated when plated on a fibronectin-coated adherent plates (stiff matrix), these mammospheres express very low levels of DRP1 compared to their parental adherent monolayer cells. Moreover, knockdown of DRP1, which typically causes mitochondrial elongation by blocking the fission process (Zhao et al., 2013, Kitamura et al., 2017), did not induce mitochondrial elongation in suspension cells. While it remains possible that DRP1 is required to generate the fragmented phenotype of mammosphere mitochondria, our results suggest that DRP1 does not play a crucial role in the regulation of mitochondrial networks once mammospheres are established. Instead, we propose that these cells modulate mitochondrial fusion to control the morphology of their mitochondria in response to ECM cues.

Cell attachment to the ECM component fibronectin stimulates integrin signaling (Buxboim et al., 2010). This signaling pathway activates a series of downstream processes that may regulate mitochondria (Sup. Figure 2A). We show that signalling events proximal to integrin activation, including FAK and Src activation, are required for mitochondrial elongation following mammosphere attachment. Further, downstream of FAK activation, the Rho GTPase Rac1 activates Arp2/3-dependent actin polymerization (Ko et al., 2014). Importantly, inhibiting Rac1 or Arp2/3 significantly disrupted mitochondrial elongation in attached mammospheres, supporting a key role for this pathway in the regulation of mitochondrial structure. Furthermore, activation of Rac in suspension cells also leads to mitochondrial elongation, demonstrating a direct link between the two processes.

Actin polymerization has previously been linked with mitochondrial fission (Moore et al., 2016), mitophagy (Onishi et al., 2021, Li et al., 2018) and mitochondrial motility (Boldogh et al., 2001). During fission, the actin-nucleating mitochondrial protein Spire1C (Manor et al., 2015), associates with ER-anchored Inverted formin 2 (INF2) (Chakrabarti et al., 2018), regulating membrane constriction and DRP1 oligomerization at the fission site. Arp2/3 complex has been also shown to influence mitochondrial division, but the mechanism is not yet defined (Li et al., 2015). Our study reveals an additional role of actin in mitochondrial dynamics, specifically in regulating mitochondrial elongation through Arp2/3-dependent actin polymerization. This is consistent with our work showing that Arp2/3-dependent actin polymerization on mitochondria is required for mitochondrial fusion (Gatti et al., 2023), and the presence of shorter mitochondria in adherent monolayer cells treated with the Arp2/3 inhibitor (Figure 5E). These different roles of actin in the regulation of mitochondria likely depend of the specific signalling pathways activated in response to distinct cellular environments. Overall, we propose here that ECM-induced mitochondrial elongation is dependent on Arp2/3-driven actin polymerisation. This alternate pathway functions independently of DRP1 activity.

In cancer cells, ECM-dependent promotion of adhesion and migration involves a metabolic shift towards increased OXPHOS and elongated mitochondria (Wu et al., 2021, Tian et al., 2019, Papalazarou et al., 2020). FAK and PAK, two Integrin-activated signaling proteins, can alter mitochondrial structure, function and energy production (Kanteti et al., 2016) (Visavadiya et al., 2016, Yang et al., 2021). Specifically, FAK inhibition results in the loss of mitochondrial membrane potential, leading to impaired mitochondrial function. Additionally, Src and FAK-activated STAT3 can directly interact with mitochondria to stimulate OXPHOS activity. (Visavadiya et al., 2016, Djeungoue-Petga et al., 2019, Guedouari et al., 2021, Guedouari et al., 2020, Lurette et al., 2022). In contrast, our results show that actin polymerization downstream of these signalling events is required to modulate mitochondrial structure in response to ECM cues in mammospheres. We thus propose that actin polymerization will have an executional role in regulating mitochondrial structure (elongation) and metabolism (OXPHOS) in response to ECM cues. This suggests that the somewhat conflicting description onthe role of the ECM and downstream signalling pathways that have been described could arise, at least in part, through differences in the actin signalling pathways activated under distinct experimental conditions. This would in turn affect mitochondrial dynamics and metabolic modulation. For instance, CSCs that require upregulation of macromolecule biosynthetic pathways during differentiation may fragment and switch to glycolysis (Serasinghe et al., 2015, Shiraishi et al., 2015), while energy demanding CSCs will gain fused mitochondria and induce OXPHOS (Rivadeneira et al., 2015).

The difference in mitochondrial usage between distinct cellular conditions could also possibly explain the discrepancy we observed between ETC components expression and OCR and ATP levels. As one possibility, we think that mammospheres could use their mitochondria to synthesize other metabolites. For example, metastatic migratory cancer cells require a high production of mitochondrial superoxide (Porporato et al., 2014), which is obtained by an exaggerated TCA cycling (Porporato et al., 2014, Porporato and Sonveaux, 2015). It is also suggested that breast cancer cells colonizing lungs utilize the proline cycle to obtain FADH2, which can be oxidized by ETC to produce mitochondrial ATP (Elia et al., 2017), or generate ATP synthesis by glycolysis and fatty acid-dependent OXPHOS, suggesting an selective rewiring of the energy substrate (Andrzejewski et al., 2017, Pascual et al., 2017).

Overall, our study reveals the complex interplay between mitochondrial dynamics and ECM-mediated actin regulation, providing insights into cellular adaptation process in changing environments. Further research is needed to comprehend actin’s role in mitochondrial function, with potential implications for understanding cancer metastasis and guiding anti-cancer therapies.

## Material and Methods

### Cell culture and transfection

MDA-MB-231 and MCF-7 cells were purchased from American Type Culture Collection. Cells in adherent monolayers were cultured in DMEM supplemented with 10% fetal bovine serum (FBS, Hyclone, UT) at 37°C in a humidified 5% CO_2_ incubator. Mammospheres were grown in DMEM supplemented with B-27 (Gibco™, 17504044), human EGF (10 µg/ml) and bovine insulin (10 µg/ml) in low-attachment plates (Nunclon sphera, Thermo Scientific™ #174932) and maintained at 37°C in a humidified 5% CO2 incubator for 3 days before harvesting. For attachment experiments, these 3-day mammospheres were spun down (1500g, 10 min) on coverslips that were previously coated with Fibronectin (Corning™ Fibronectin, Human Fischer Scientific, #CB-40008A) or Poly D Lysine (Sigma-Aldrich #P6403) where indicated. Cells were then kept at 37°C in a humidified 5% CO_2_ incubator until they were fixed. The same procedure was followed for western blot except that cells were spun down on standard cell culture dishes for adherent cells.

### siRNA treatment

MDA-MB-231 cells were seeded onto 24-well plates and transfected with 15nM of DRP1 siRNA (Thermo Fisher Scientific, Silencer Select, #4390771) or negative siRNA (Thermo Fisher Scientific, Silencer Select, #4390843) using siLenFect lipid reagent (Bio-Rad, #1703361). After 24 hrs, cells were collected for either western blotting or seeded onto coverslips for immunofluorescence.

### Manipulation of actin polymerization

The following proteins were manipulated using chemical inhibitors/activators: F-Actin depolymerization, Cytochalasin D (0.1 μg/ml Sigma Aldrich #C8273); FAK, small-molecule inhibitor PF-573228 (10 μM; Sigma Aldrich #PZ0117); Src kinase, Src Inhibitor-1 (10 μM; Sigma Aldrich #S2075); Rho GTPase, Rho Inhibitor I (C3 Transferase from *Clostridium botulinum* covalently linked to a cell penetrating moiety; 10μM; Cytoskeleton, Inc #CT04-A); Rac GTPase, Rac Inhibitor III (2 μM; MilliporeSigma™ #55351310MG); Arp 2/3 complex, CK-666 (10 μM; Sigma Aldrich #182515); Formin, FH2 domain inhibitor SMIFH2 (2µM; Sigma-Aldrich #344092); Rac activation, Rho/Rac/Cdc42 Activator I (10 μM; Cytoskeleton, Inc #CN04-A). All treatments were applied for a duration of 1 hour prior to fixation with 4% PFA, followed by subsequent immunofluorescence and confocal imaging.

### Immunofluorescence

Cells were plated on glass coverslips (Fischer Scientific #12541000CA) and allowed to adhere overnight to establish monolayer cultures. Mammospheres, on the other hand, were anchored to coverslips through centrifugation at 3,000 rpm for 10 minutes, followed by incubation for a period ranging from 1 to 6 hours. Mammospheres in suspension were collected by centrifugation in a microtube. The cells were then fixed with 4% paraformaldehyde for 15 minutes at room temperature (RT). Cells were then permeabilized with 0.2% Triton X-100 in PBS and blocked with 1% BSA / 0.1% Triton X-100 in PBS. Cells were then incubated with a primary antibody against TOM20 (Rb,1:250; Abcam #ab186735,) followed by fluorescent-tagged secondary antibodies (1:500; Jackson Immunoresearch). Cells were co-stained with Rhodamine-phalloidin (1:250; Sigma Aldrich #P1951), and DAPI (1:100; Invitrogen, Thermo Fisher #D1306).

### Image processing and analysis

The images were acquired using a Leica TSC SP8 confocal microscope with a 63×/1.40 oil objective using the optimal resolution for the wavelength (determined using the Leica software). All image manipulation and analysis were done in Image J/Fiji. The images shown are from single focal planes unless stated otherwise. For mitochondrial length analysis, a cell was binned in a specific category if at least 70% of mitochondria within that cell had the following length: less than 0.2 µm for fragmented, between 0.2 and 0.6µm for intermediate and more than 0.6 µm for elongated. Cells where mitochondrial networks were not clearly identifiable were excluded from the analysis.

### Analysis of OPA1 oligomers

OPA1 oligomerization within intact cells was done as described (Patten et al., 2014). Briefly, cells were treated with the cell-permeable cross-linking agent BMH (Thermo Scientific) at a concentration of 1 mM for 20 minutes at 37°C. Following the crosslinking procedure, BMH was quenched by rinsing twice using PBS containing 0.1% beta-mercaptoethanol (BME). Subsequently, the cells were collected and lysed in 10 mM Tris-HCl at pH 7.4, 1 mM EDTA, 150 mM NaCl, and 1% Triton X-100 with 0.1% BME, and the lysate was subjected to Western blot on NuPAGE Novex 3–8% Tris-acetate gradient gels (Invitrogen™ # EA0375BOX), transferred to a nitrocellulose membrane and blotted against OPA1 (Anti-Mouse, BD Biosciences, #612606) and mtHsp70 (Anti-Mouse, Clone: JG1, Invitrogen MA3028) antibody.

### Western blots

Cells were lysed in 10 mM Tris-HCl, pH 7.4, 1mM EDTA, 150 mM NaCl, 1% Triton X-100, complemented with a protease inhibitor cocktail (Sigma-Aldrich #11836170001) and phosphatase inhibitors (Sigma-Aldrich), kept on ice for 10 min and centrifuged at 16,000 x g for 10 minutes. Protein supernatants were collected, and protein concentration was estimated by DC protein assay (BioRad). For SDS-PAGE, 30 µg of proteins were mixed with 1X Lammeli buffer containing β-mercaptoethanol, then subjected to SDS-PAGE, transferred to a nitrocellulose membrane and blotted with the indicated antibodies (DRP1 (Anti-Mouse, 1:1000; BD Transduction Laboratories, #611112), Phospho-DRP1 (Ser616) (Anti-Rabbit, 1:1000; Cell signaling technology, (Clone D9A1) #4494S), MFN1 (Anti-Rabbit, 1:1000; Abcam, [EPR21953-74] #ab221661), OPA1(Anti-Mouse, BD Biosciences, #612606), NDUFA9(Anti-Rabbit, 1:1000; Abcam #ab128744), OSCP (Anti-Mouse, 1:1000; Santa Cruz Biotechnology (clone A-8) #sc-365162) UQCRC2 ((Anti-Mouse, 1:1000; Santa Cruz Biotechnology (clone G-10) #sc-390378) Phospho-FAK^Y397^ (Anti-Rabbit, 1:1000; ; Invitrogen, (clone 31H5L17) # 700255), FAK(Anti-Rabbit, 1:1000; ) TOM20 (Anti-Rabbit, 1:1000; Abcam, ab186735), SOX-2 (Anti-Rabbit, 1:1000 #AB5603 ), ALDH1A1 (Anti-Mouse, 1:1000, # SC-166362), HRP-tagged Actin (1:10,000). Membranes were then incubated with horseradish peroxidase-conjugated secondary antibodies (1:5000; Jackson Immunoresearch) and visualized by enhanced chemiluminescence (Thermo Fisher scientific) using a Bio-Rad imaging system.

### OCR measurements

Oxygen Consumption Rates (OCR) were measured using the Seahorse XF Cell Mito Stress Test Kit (Agilent, Cat. 103708-100) with a Seahorse XFe96 analyzer (Agilent), following the instructions provided by the manufacturer. For mammospheres attached to fibronectin, a total of 40,000 cells from the 3D culture were placed onto a Seahorse XF96 culture plate coated with fibronectin (10 µg/ml), in DMEM media supplemented with hEGF (10 ng/ml), insulin (10 µg/ml), and B-27 supplement(Dey et al., 2009). The cells were incubated for 6 hours at 37°C, 10% CO<u>_2_</u> promoting attachment of mammospheres. Following this, a transition to XF DMEM medium was carried out. Meanwhile, 40,000 AM and MS cells were seeded onto Seahorse XF96 plates in XF DMEM, supplemented as indicated by the manufacturer. AM cells were seeded and allowed to attach for 6 hours while MS were transferred at the time of the assay. The cells were then incubated at 37 °C without CO_2_ for 1 hour, before proceeding immediately to perform OCR and ECAR analyses. OCR was monitored by sequentially introducing oligomycin (1 μM), FCCP (0.25 μM), and rotenone/antimycin A (0.5 μM) and non-mitochondrial respiration was subtracted for the calculation of mitochondrial respiration. Basal OCR and ATP-linked OCR were calculated according to manufacturer instructions.

### ATP and Lactate assay

ATP assays were performed using CellTiter-Glo^®^ Luminescent Cell Viability Assay Kit (Promega). Cells were seeded at a density of 20,000 cells per well in 96-well plates (transparent and opaque walled plates) and incubated overnight at 37°C. For attached mammospheres, cells in suspension culture were seeded onto fibronectin-coated wells six hours prior to performing the assay. Where indicated, the cells in monolayer, mammospheres and mammosphere attached condition were then treated with Oligomycin (10 µM) for 1 hour before the end of the incubation period. CellTiter-Glo^®^ Reagent was added to the cells in the opaque plates at a 1:1 ratio and incubated at room temperature for 10 minutes in a dark environment. The luminescence emitted was then measured at a 100% gain value in a plate reader.

The transparent plate was used for both lactate and protein estimation. For lactate estimation, a portion of the medium (10 μl) from each well was transferred to fresh wells in duplicate and the volume adjusted to 50 µl with PBS. Subsequently, 50 μl of Lactate reagent from Promega (# J5021) was added and incubated for 10 minutes at room temperature. The resulting absorbance was recorded at a wavelength of 450 nm in a plate reader. The media from the transparent plates was removed leaving the cells at the bottom. The cells were rinsed with PBS and subsequently lysed (10 mM Tris-HCl (pH 7.4), 1mM EDTA, 150 mM NaCl, and 1% Triton X-100). The protein concentration in this lysate was estimated using the DC protein assay (BioRad) and used to normalize ATP and lactate values.

### TMRM experiments

Mitochondrial membrane potential was evaluated using the potentiometric dye tetramethyl rhodamine methyl ester (TMRM; Sigma Aldrich) at a final concentration of 10 nm. For the assay, 3 x 10^5^ cells/well of both adherent and suspension cells were seeded in a 6-well microplate and left overnight in an incubator (5% CO_2_, 37°C). Each experiment was performed in duplicate. The cells were then incubated 30 minutes with TMRM (10 nm) in DMEM. Note: for suspension cells, mammospheres were trypsinized to obtain single cells before TMRM treatment. Adherent cells were then washed with PBS, trypsinized, and brought to a single cell suspension. Suspension cells were collected and washed with PBS. The blank control cells untreated with TMRM were also collected. The cell pellets were then resuspended in PBS and analyzed using flow cytometry. The TMRM fluorescence was examined using a Cytoflex FACS analyzer (Beckman) with excitation at 514 nm and detection at 570 nm.

### Animals

All animal experiments adhered to the principles outlined in the guidelines of laboratory animal care (NIH publication no. 85-23, revised in 1985) and were conducted with the approval of the Institutional Animal Ethical Committee, Government of India (Registration Number 885/ac/05/CPCSEA). The experimental procedures strictly followed the regulations specified in the Indian Laws of Animal Protection (ILAP). Female BALB/c mice, weighing 20 ± 2g, were obtained and housed under standard laboratory conditions, maintaining a temperature of 25 ± 2°C, relative humidity of 50 ± 15%, and a 12-hour light-dark cycle throughout the experiment. The mice were provided with food pellets, and water was given *ad libitum* to facilitate acclimatization to the laboratory environment. (Elia et al., 2017).

### Mice mammary tumor development

Mammary tumors were developed in female BALB/c mice (n=3). Viable triple negative mouse mammary carcinoma cells 4T1 (10^5^ cells) suspended in 200 µl of PBS were injected into the inguinal 4th mammary fat pad of mice. Tumors were allowed to develop for 14 days. Normal mice received only 200 µl of PBS. Eventually, the normal mammary gland and the mice-mammary tumors were excised from the control and mammary tumor-bearing animals respectively. The tissues were then processed for subsequent experiments (Elia et al., 2017).

### Data analysis and statistics

All graphs and statistical analysis were done using R. Immunofluorescence data were quantified and images representative of at least three independent experiments shown (exact “n” are in the quantification figures). Data is represented as average ± SD as specified in figure legends. Statistical significance was determined using Student’s t test (between 2 groups) or one-way ANOVA with a Tukey post hoc test (multiple comparisons).

### Author contributions

Conceptualization, PG, PM, UC, and MG; Methodology, PG, PM, PDT, WF, LCC, UC and MG; Investigation, PG, PM, PDT, JK, UC, and MG; Resources, LCC; Writing–Original Draft, PG and MG; Writing – Review and Editing, all the authors; Supervision, LCC, UC, and MG; Funding Acquisition, MG

## Acknowledgements.

This work was supported by a grant from the Natural Sciences and Engineering Research Council of Canada to MG, a team grant from Fonds de Recherche du Québec-Nature et Technologies (#2024-PR-326143) to M.G and L.C.C. and a Department of Science, Technology and Biotechnology, Government of West Bengal grant to UC (Sanction No.: 140 (Sanc.)-BT/P/Budget/RD-75/2017 dated 16.11.2018). PG was supported by a Queen Elizabeth II Diamond Jubilee Scholarship and an FRQ-NT scholarship, PDT was supported by a fellowship from the Council for Scientific and Industrial Research, Government of India (File No.-09/028(1066)/2018-EMR-I). PG received a Université du Québec à Trois-Rivières Programme d’aide à l’internationalisation de la science scholarship. W.F received PhD fellowships from the Armand-Frappier Foundation and Fonds de la Recherche du Québec-Santé (FRQS). L.C.C is receiving a research scholar (Senior) salary support from FRQS.

---

**Supplemental Figure 1.**
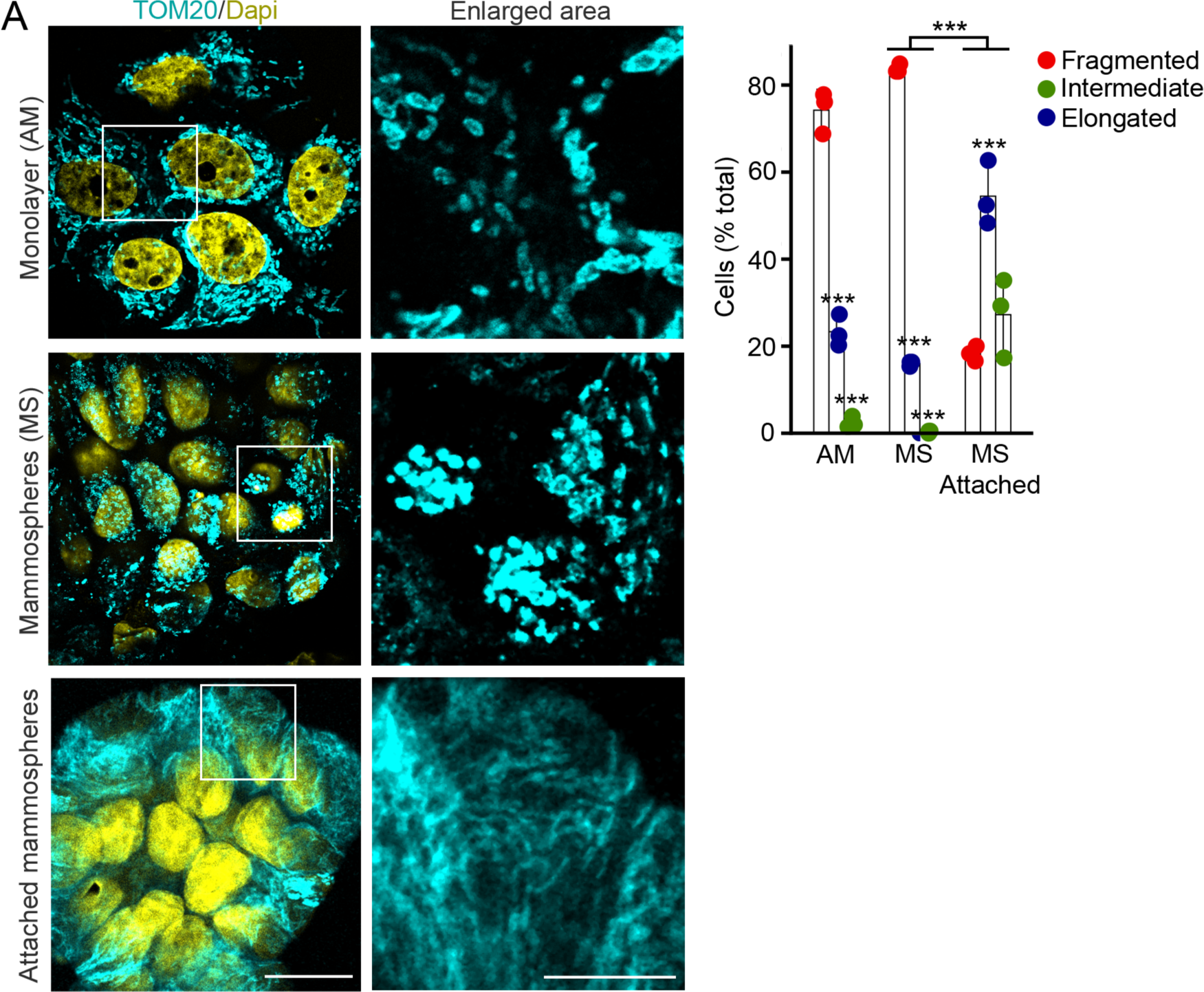
Mitochondrial fragmentation in MCF7 mammospheres in suspension. (A) MCF7 were grown as AM, MS or MS attached for 6 hours on glass coverslips, and their mitochondria were marked with an antibody against TOM20 (mitochondria, cyan; with nuclei stained with DAPI, yellow). (B) Representative images. Scale bar 10 µm. (C) Quantification of 3 independent experiments, with each point represents an individual experiment. Bars show the average ± SD. *** p<0.001. Two-way ANOVA.

**Supplemental Figure 2.**
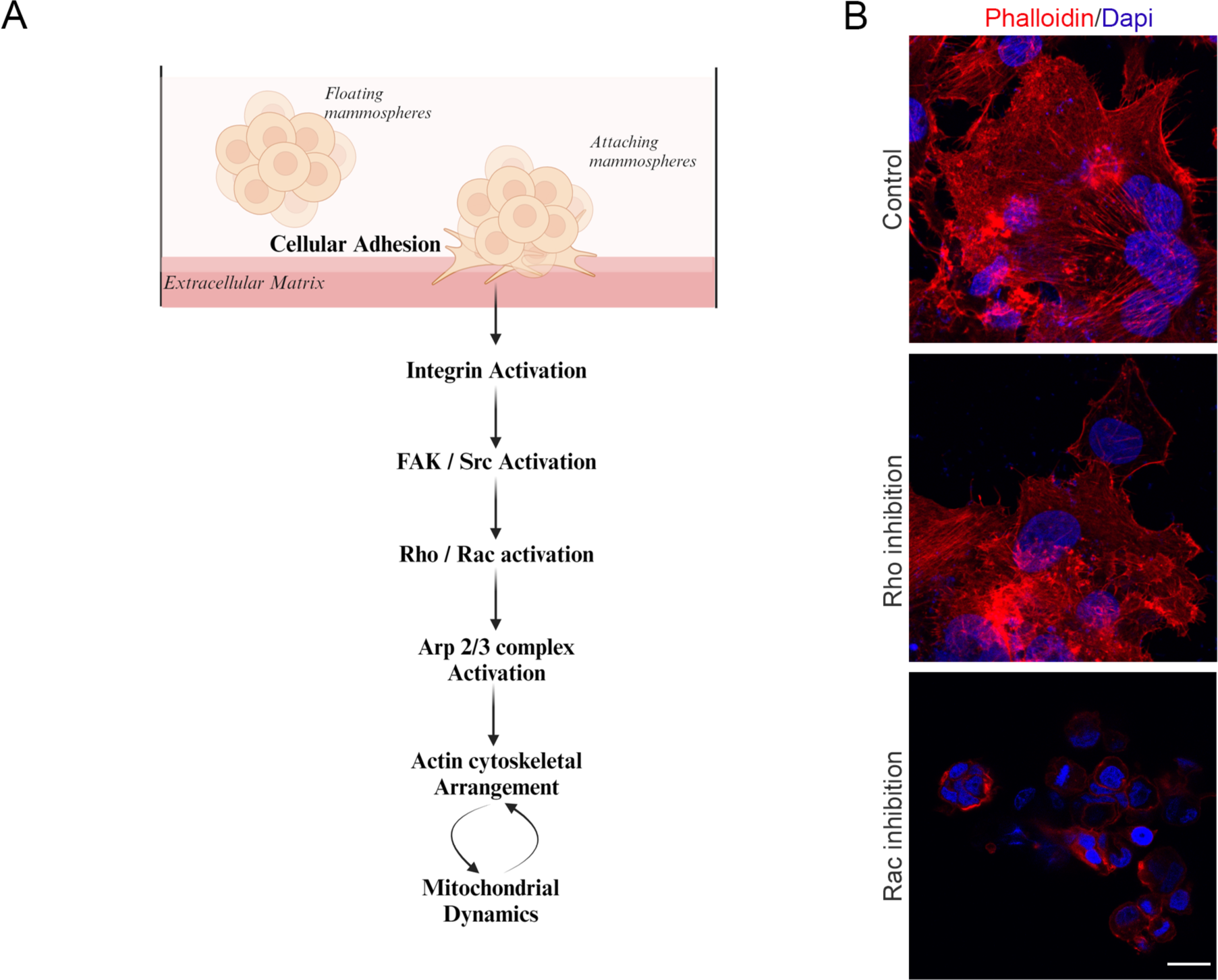
Mitochondrial fragmentation in attached MS upon Rac inhibition. (A) Model for the ECM-induced cytoskeletal pathway regulating mitochondrial dynamics in mammospheres. (B) Representative images of mammospheres attached to fibronectin for 1 hour in the absence or the presence of the Rac1 (Rac Inhibitor III; 2 μM) and Rho (Rho Inhibitor I, 10μM) inhibitors and marked for Actin (Phalloidin, Red) and nuclei (Dapi, Blue). Scale bar 10 µm.

